# Lack of Host Reproductive Trade-offs Associated with Fecundity Compensation Response to Parasitic Castration

**DOI:** 10.1101/2021.06.28.450230

**Authors:** Annabell A. Davis, J. Trevor Vannatta, Stephanie O. Gutierrez, Dennis J. Minchella

**Affiliations:** Department of Biological Sciences, Purdue University, 915 West State Street, West Lafayette, Indiana, USA

**Keywords:** *Schistosoma mansoni*, *Biomphalaria glabrata*, Life history, Terminal investment, Trade-offs, Resource allocation

## Abstract

Host-parasite coevolution may result in life-history changes in hosts that can limit the detrimental effects of parasitism. Fecundity compensation is one such life-history response, occurring when hosts increase their current reproductive output to make up for expected losses in future reproduction due to parasitic infection. However, the potential trade-offs between quantity and quality of offspring produced during fecundity compensation are relatively unexplored. This study uses the trematode, *Schistosoma mansoni*, and its snail intermediate host, *Biomphalaria glabrata*, to better understand the impacts of this host life-history response. Measures of host reproductive output as well as offspring hatching success and survival were collected to assess the reproductive consequences of infection. Infected snails exhibited fecundity compensation (increase in the number of eggs laid compared to controls) and had a higher probability of laying any eggs at all. Infection status did not play a significant role in hatching or offspring survival to maturity. However, the age of the parental snail had a significant impact on hatching success, as offspring from older parents demonstrated a higher hatching success rate. Overall, the lack of an apparent trade-off between quantity and quality of offspring suggests that infected parental snails invest more resources towards reproduction in order to maintain the fitness of their offspring, possibly at the expense of their own longevity.

## Introduction

The intimate association between hosts and parasites inevitably shapes their evolutionary paths. Parasites evolve to maximize their fitness using the energy and resources of its host, while the host simultaneously evolves to mitigate these adverse effects (Sorensen & Minchella, 2001). To diminish the negative fitness impacts of parasitic infections, changes in host life-history traits can evolve (Minchella, 1985). Parasites can utilize castration to consume the resources their hosts allocate for reproduction towards their own fitness instead (Lafferty & Kuris, 2009). In response to the early stages of parasitic castration and threat to survival, hosts can increase their current reproductive output to make up for lost future reproduction. This life-history response is termed ‘fecundity compensation’ (Minchella & LoVerde, 1981) or ‘terminal investment’ (Duffield et al., 2017).

Fecundity compensation due to parasitism has been documented in a variety of organisms including birds, mammals, insects, molluscs, and crustaceans (Duffield et al., 2017). For example, the house martin bird, *Delichon urbica* is infected by both *Haemoproteus* or *Plasmodium* species, resulting in increased mortality but also increased reproductive investment and success by laying larger clutches and laying earlier (Marzal et al., 2008). The fruit fly, *Drosophila nigrospiracula*, can be infected by an ectoparasitic mite, *Macrocheles subbadius*, which reduces the life span of the host. However, when infected, males increase their courtship activity in an effort to compensate for lost future reproduction opportunities (Polak & Stamer, 1998). Thus, fecundity compensation is a widespread phenomenon and occurs not only in response to parasitic infection, but also to a variety of factors that can negatively impact future reproduction including age, nutrition-dependent conditions, and predation (Duffield et al., 2017).

While fecundity compensation has been shown to occur as a life history response to parasitism, it is unclear whether there is a trade-off between the quantity and quality of host offspring. That is, do eggs produced during fecundity compensation have the same hatching success and survival to maturity compared to eggs produced by uninfected individuals? In the parasitoid wasp *Lysiphlebus orientalis* and host soybean aphid *Aphis glycines* system, infection results in decreased overall fecundity but larger embryos, possibly accounting for the increased reproductive rate seen in those offspring (Kaiser & Heimpel, 2016). In deer mice, *Peromyscus maniculatus*, infected by the trematode, *Schistosomatium douthitti*, there is no change in the number of offspring produced, but offspring from infected individuals are born and weaned at heavier masses. This increase in available resources for the offspring can increase their survival and increase their reproduction potential once they reach maturity (Schwanz, 2008)

In the current study, *Schistosoma mansoni* and its intermediate host, the freshwater snail *Biomphalaria glabrata*,were utilized to investigate the consequences of parasitic infection on host reproduction and progeny success. *S. mansoni* is the etiologic agent for the human parasitic disease schistosomiasis, which affects over 230 million people worldwide. Freshwater snails of the genus *Biomphalaria* are infected by *S. mansoni* miracidia that hatch from eggs in human faeces. Once infected, snails release *S. mansoni* cercariae into the water where they can infect humans and the cycle repeats (CDC, 2018).

*B. glabrata* snails are eventually castrated by *S. mansoni*; however, soon after exposure, there is an increase in the rate of host reproductive output (fecundity compensation; Minchella & LoVerde 1981). This project aims to evince the eventual quality of the offspring produced by infected snails in order to better understand their coevolutionary interactions. For many species, there is a trade-off between size and number of offspring (Smith & Fretwell, 1974). Furthermore, consistent positive directional selection for larger body size has been widely documented, indicating an association between body size and fitness (Kingsolver et al., 2012). Therefore, it is predicted that the increase in quantity of eggs produced during the fecundity compensation period will reduce the quality of the individual offspring.

## Material and methods

Ninety M line *Biomphalaria glabrata* snails between 8-10mm were randomly assigned to two groups. Fifty snails were exposed to NMRI *Schistosoma mansoni* while 40 served as unexposed controls. For *S. mansoni* exposure, infected mice were euthanized in accordance with Purdue Animal Care and Use Committee Protocol # 1111000225 to isolate *Schistosoma* eggs. The collected eggs were placed in freshwater for approximately 60 minutes to allow the miracidia (the infective stage for the snails) to hatch. The resulting miracidia were transferred to 6-well plates containing snails. Each snail was exposed individually using five miracidia or sham exposed (unexposed group) for 24 hours.

Snails were kept in individual jars with a piece of Styrofoam for laying eggs and were fed lettuce *ad libitum*. Beginning at 4 weeks post-exposure, infection status was checked weekly by placing snails under fluorescent light for one hour and checking for the presence of cercariae. Snails that released cercariae were designated as infected. Snails that were exposed to miracidia but did not release cercariae were designated as exposed but uninfected.

The reproductive output and accuracy in laying eggs was compared between infected and uninfected snail parents as well as hatching success and successful survival to maturity of their offspring. Egg laying accuracy was considered the ability to allot a single yolk to a single egg case. Mistakes that were seen were empty egg cases and multiple yolks in one egg. To determine reproductive output, each week post exposure the number of egg masses laid by each parent was counted as well as the number of eggs in each egg mass. This was achieved by carefully removing the egg masses from the Styrofoam and placing them on glass slides. Under a light microscope, the number of eggs were counted as well as the number of eggs with and without yolks.

In addition, each week (for a total of 5 weeks or until parasitic castration of the infected parent snail), approximately 10 and 20 eggs with yolk were retrieved from each parent and used for assessing hatching success and survival to maturity, respectively. Hatching success was determined by placing approximately 10 eggs in a well plate and counting the number of snails that hatched from the egg mass within two weeks of being laid. Survival to maturity was determined by the proportion of snails that grew to 9 mm in diameter out of the original clutch (Théron et al., 1998). This was done by placing approximately 20 eggs in individual jars and allowing them to hatch and mature. Snail offspring were fed lettuce *ad libitum*. Each week, snails were measured using a calliper and any snails 9 mm or larger were recorded and removed from the jars.

All statistics were done using R version 3.6.3 (R Core Team, 2020) and all pairwise comparisons were done using the multivariate t distribution p value correction in the emmeans package (Lenth, 2021). For analysis of fecundity compensation, we performed a zero-inflated mixed effects model with negative binomial error distribution in the glmmTMB package (Brooks et al., 2017). This was due to the abundance of zeros in our data and data overdispersion. Weekly egg laying data from uninfected and infected snails was compared over the first three weeks after *S. mansoni* exposure (the typical fecundity compensation window) with parent snail as a random factor.

To determine the impact of infection status compared to parent age, we ran our models two ways using the lme4 package (Bates, et al., 2015). First, hatching success and survival to maturity of offspring was analysed using a generalized linear mixed model with binomial error distribution with treatment group as a fixed factor and parent snail, maturation jar, and week laid (parent age) included as random factors. Second, hatching and survival to maturity were analysed with the week the parents laid the eggs, indicating parent age, as a fixed effect with parent snail, maturation jar, and infection status as random factors.

Egg laying accuracy was analysed for infected, uninfected, and exposed but uninfected parent snails using a binomial GLMM with parent snail and week included as random factors.

## Results

### Fecundity compensation

We observed fecundity compensation in *S. mansoni* infected *B. glabrata* snails occurring in two forms. First, infected snails had a higher probability of laying an egg mass than uninfected snails, especially in weeks 2 and 3 (Table 1 and Figure 1a). This difference is visible in Figure 1, where the narrower base (fewer 0 values) indicates a higher probability of laying eggs for the infected group. In addition, there was a significant difference in the number of egg masses laid, where infected snails laid more egg masses, especially in earlier weeks (Table 1 and Figure 1a). Very similar results were seen for the number of eggs laid by snails (Table 2 and Figure 1b).

**Table 1:** Mixed effects model results with negative binomial error distribution for egg masses laid. Infected snails are compared relative to uninfected (intercept) with parent snail as a random factor. The estimate of the zero-inflation model represents the probability of obtaining a zero value, or no laying.

| **Conditional Model** | **Estimate** | **Standard Error** | **Z Value** | **P Value** |
| --- | --- | --- | --- | --- |
| Intercept | 1.40985 | 0.14660 | 9.617 | **< 0.0001** |
| *S. mansoni* Infected | 0.44550 | 0.19515 | 2.283 | **0.0224** |
| Week | 0.16006 | 0.06115 | 2.618 | **0.0089** |
| *S. mansoni* Infected by Week interaction | -0.16907 | 0.08301 | -2.037 | **0.0417** |
| **Zero Inflation Model** | **Estimate** | **Standard Error** | **Z Value** | **P Value** |
| Intercept | -6.753 | 1.997 | -3.382 | **0.0007** |
| *S. mansoni* Infected | 6.617 | 3.226 | 2.051 | **0.0402** |
| Week | -2.664 | 1.102 | -2.417 | **0.0156** |
| *S. mansoni* Infected by Week interaction | -6.494 | 2.294 | -2.830 | **0.0047** |

**Table 2:** Mixed effects model results with negative binomial error distribution for eggs laid. Infected snails are compared relative to uninfected (intercept) with parent snail as a random factor. The estimate of the zero-inflation model represents the probability of obtaining a zero value, or no laying.

| **Conditional Model** | Estimate | Standard Error | Z Value | P Value |
| --- | --- | --- | --- | --- |
| Intercept | 3.30809 | 0.19374 | 17.075 | **< 0.0001** |
| *S. mansoni* Infected | 0.55269 | 0.25606 | 2.158 | **0.0309** |
| Week | 0.30759 | 0.7076 | 4.347 | **< 0.0001** |
| *S. mansoni* Infected by Week interaction | -0.32351 | 0.09619 | -2.417 | **0.0156** |
| **Zero Inflation Model** | Estimate | Standard Error | Z Value | P Value |
| Intercept | -7.3657 | 1.8354 | 4.013 | **< 0.0001** |
| *S. mansoni* Infected | 6.9200 | 3.814 | 2.246 | **0.0247** |
| Week | -1.6196 | 0.6626 | -2.444 | **0.0145** |
| *S. mansoni* Infected by Week interaction | -6.494 | 2.294 | -2.830 | **0.00465** |

**Figure 1:**
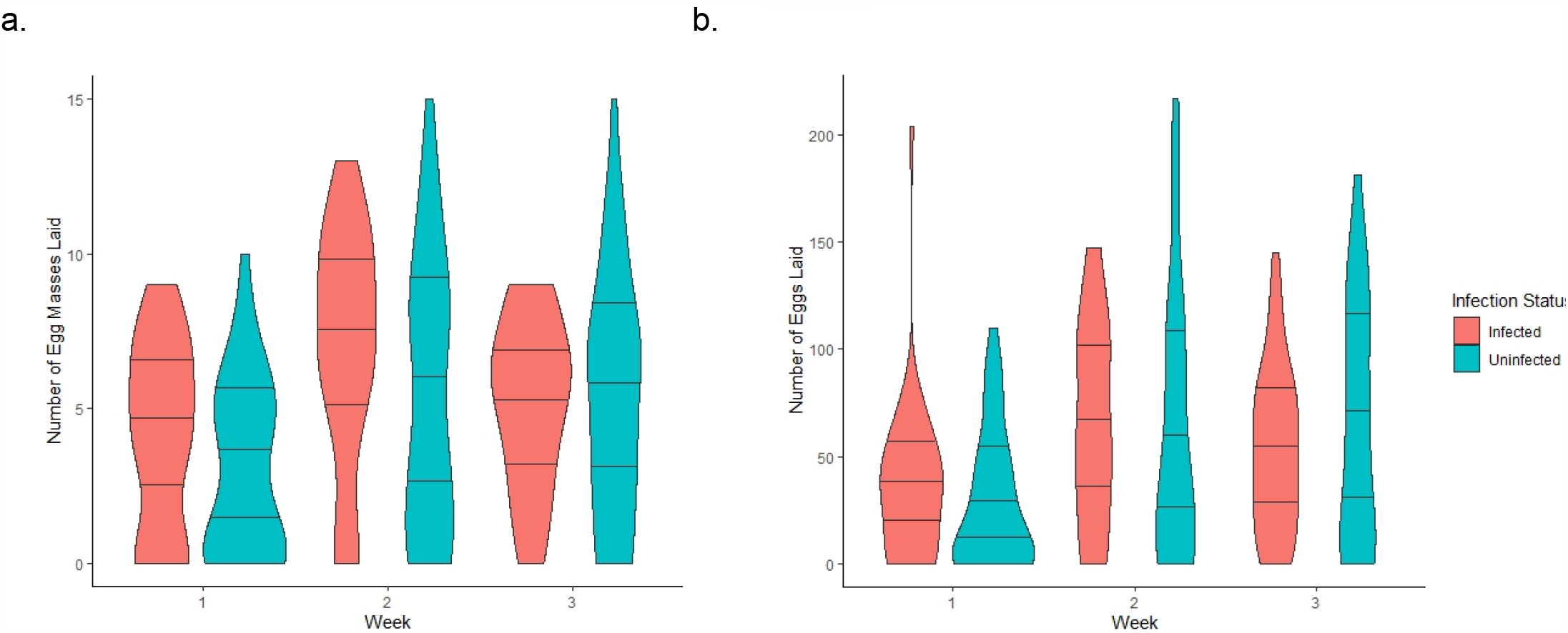
Violin plot of the number of egg masses laid (a) and number of eggs laid (b) by *B. glabrata* snails either uninfected or infected with *S. mansoni* each week after exposure until week 3 when castration begins for infected snails. Lines within the violin plots delineate data quantiles.

### Hatching success

We saw that infection status was not a significant predictor of *B. glabrata* offspring hatching success (p=0.232). However, week the parents laid the eggs, indicating parent age, was a significant predictor of hatching success (p≤0.001, Table 3). Eggs that were laid later, or older parents, had a greater proportion of offspring hatch (Figure 2).

**Table 3:** Generalized linear mixed model results comparing weeks post experiment start (parent snail age) and likelihood offspring hatch with parent snail, maturation jar, and infection status as random factors.

|  | Estimate | Standard Error | Z Value | P Value |
| --- | --- | --- | --- | --- |
| Intercept | 1.5542 | 0.3909 | 3.977 | < **0.0001** |
| Week (age) | 0.4238 | 0.1271 | 3.335 | **0.0009** |

**Figure 2:**
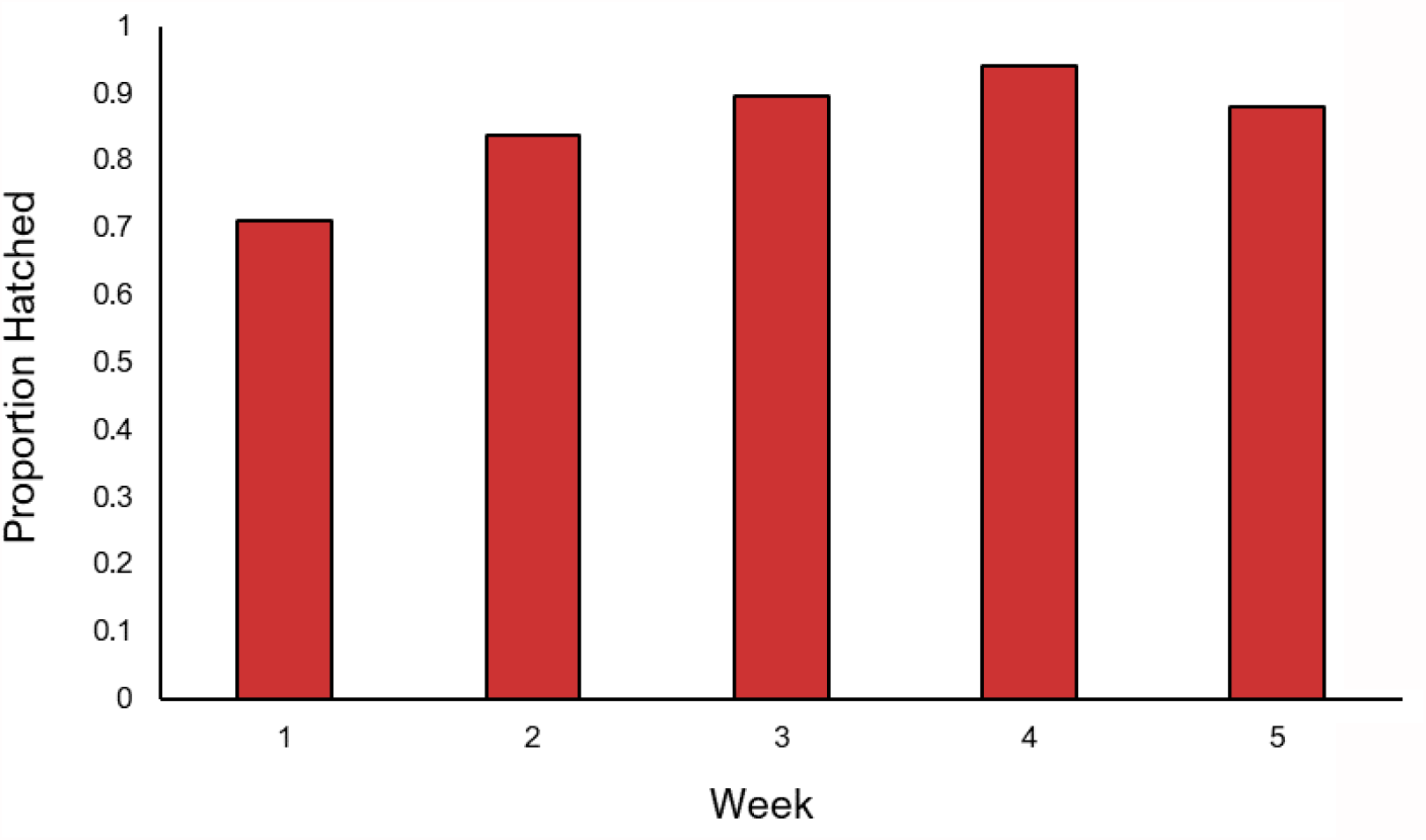
Week post experiment start (parent snail age) and the proportion of *B. glabrata* offspring hatched.

### Survival to maturity

Infection status played no significant role in the chances *B. glabrata* offspring surviving to maturity (p=0.727). There was also no effect of the week eggs were laid (parent snail age) on survival to maturity (p=0.459).

### Egg laying accuracy

*B. glabrata* snails exposed to *S. mansoni*, but not infected, were more likely to make errors in allotting a single yolk to a single egg case than either infected (p=0.0249, Table 4) or uninfected snails (p=0.0285, Table 4) as seen in Figure 3.

**Table 4:** Pairwise comparison results of infected, exposed but uninfected, and uninfected snails using generalized linear mixed model with binomial error distribution with snail and week as random factors.

| Contrast | Estimate | Standard Error | Degrees of Freedom | Z Ratio | P Value |
| --- | --- | --- | --- | --- | --- |
| Exposed but<br>Uninfected<br>versus<br>Uninfected | -0.9425 | 0.371 | Inf | -2.541 | **0.0285** |
| Uninfected<br>versus<br>Infected | 0.0221 | 0.259 | Inf | 0.086 | 0.9959 |
| Exposed but<br>Uninfected<br>versus<br>Infected | 0.9646 | 0.372 | Inf | 2.592 | **0.0249** |

**Figure 3:**
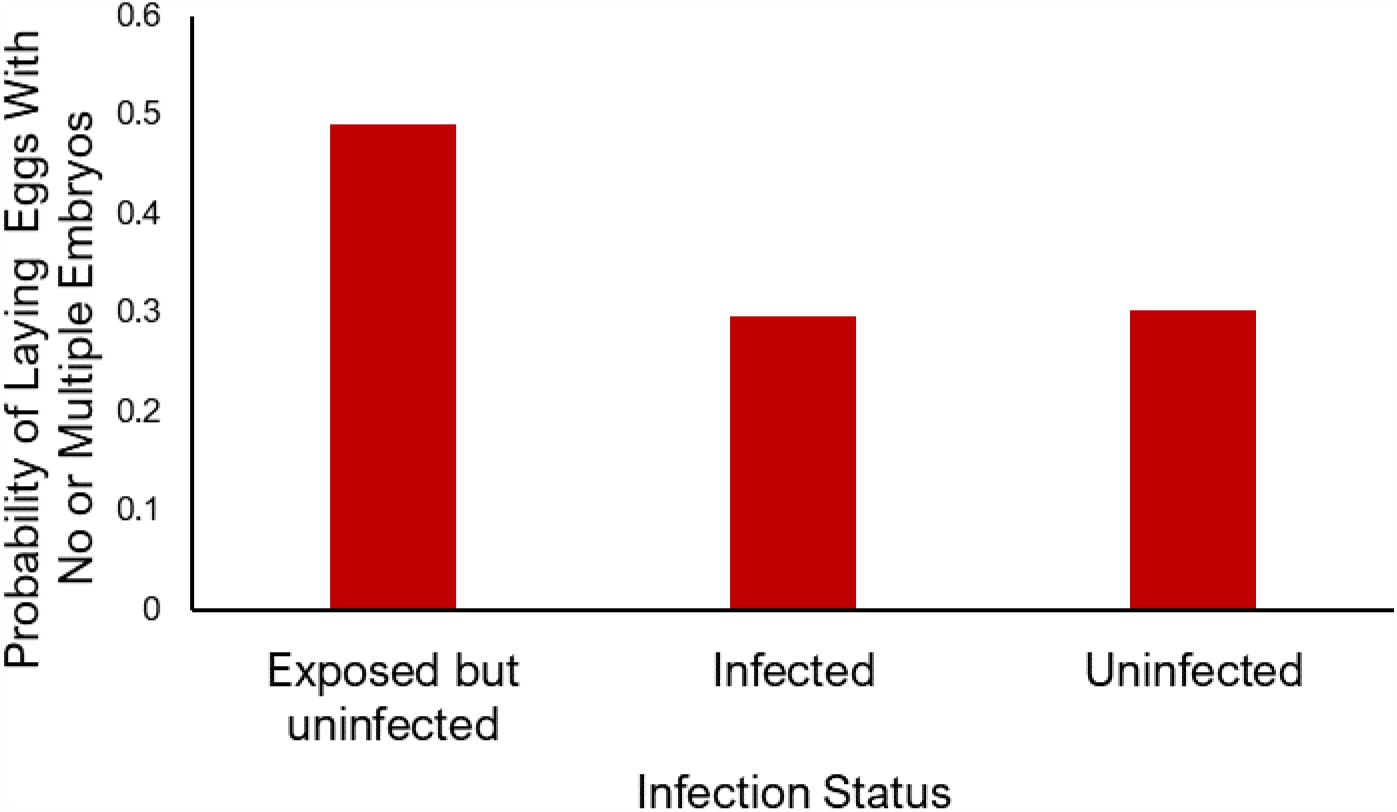
Infection status of *B. glabrata* and the probability of laying eggs with no or multiple embryos in a week.

## Discussion

The interactions between host and parasites result in reciprocal selective pressures, leading to coevolutionary arms races (Minchella, 1985). Fecundity compensation is a host life history response to parasitic castration (Minchella and LoVerde, 1981). While fecundity compensation is widely documented, the potential tradeoff between the increased number of offspring and the quality of each offspring is relatively unexplored. In this study, several reproductive measures including reproductive output, ability to allot a single yolk to an egg, hatching success of offspring, and successful growth of offspring to maturity were collected to compare offspring produced during fecundity compensation with those from uninfected snails.

We observed fecundity compensation in infected *B. glabrata* in two forms. In earlier weeks, infected and uninfected snails had similar probability of laying eggs. However, when infected snails did lay, they laid more egg masses and eggs. In later weeks, infected snails have a higher probability of laying eggs, but the number of egg masses and eggs laid is similar to uninfected snails. To the best of our knowledge, fecundity compensation has only referred to the increased number of offspring produced. This study demonstrated that fecundity compensation can also be viewed as the increased likelihood to produce offspring at all.

The difference in the forms of fecundity compensation may relate to different stages of castration as parasitic castration can be a gradual process (Lafferty & Kuris, 2009). In this study, when infected snails are early in the castration process, they produce more egg masses and eggs when they did lay. Later in the castration process, as the number of egg masses and eggs laid levels out, but the likelihood of laying increases. This adaptation may allow the snail host to utilize its reproductive system to the best of its ability (under challenging circumstances) to produce as many offspring as possible for the next generation.

We predicted a trade-off between offspring quality and quantity of infected versus uninfected snails during fecundity compensation. However, our results demonstrate that infected snail offspring have the same ability to hatch and reach adulthood as uninfected snail offspring. We also examined the ability of parent snails to allot a single yolk to a single egg, representing the accuracy in laying eggs and saw no difference between infected and uninfected snails. In life history theory, resources are limiting, and must be budgeted appropriately to remain evolutionarily advantageous (Stearns, 1992). Resources can be allocated towards the short-term growth of an individual, reproduction, or long-term health and survival (Sorensen & Minchella, 2001). The lack of trade-offs between offspring quality and quantity in infected versus uninfected snails in this study implies that exposed snails can allocate enough energy and resources to reproduction to lay healthy eggs that produce viable offspring. However, without an increase in energy/resource intake, increased investment in reproduction must diminish another life history trait. Fecundity compensation may play a role in the decrease in snail longevity seen during *S. mansoni* infection (Sturrock & Sturrock, 1970). In our study, snails were fed ad libitum. Thus, future studies should examine if these results are consistent in resource limited environments.

Additionally, the later eggs were laid, indicating older parents, the more likely they were to hatch. The association between parental age/size and reproductive performance has been seen in multiple studies with a variety of iteroparous animals (Fox & Czesak, 2000; Marshall et al., 2008; Sakai & Harada, 2001; Yanagi & Takahisa, 2002). In general, older parents lead to offspring of greater fitness which can be measured in a variety of ways (Marshall et al. 2010). This includes hatching success as seen with apple snail (*Pomacea canaliculata*), Japanese Quail (*Coturnix japonica*), Western Gull (*Larus occidentalis*), and Ring-billed Gull (*Larus delawarensis*) (Estoy et al., 2002; Haymes & Blokpoel, 1980; Seker et al., 2004; Sydeman et al., 1991). In snails, this pattern could potentially be explained evolutionarily by energy or resource allocation differences between different ages and stages of development (Gérard & Théron, 1997). According to the dynamic energy budget model of Kooijman (1993), younger parents may allocate more resources towards growth and development, but as they become more mature, they shift allocation more towards reproduction.

Our results were limited to only a portion of the entire lifespan of *B. glabrata* parents. In future studies, this can be explored further by examining the entire lifespan of *B. glabrata* parents to find the overall pattern and optimal parental age for offspring fitness. Offspring fitness can be examined using many other parameters, such as size of offspring and offspring life-history traits such as longevity (Plaistow et al., 2015).

We also observed that snails that were exposed to *S. mansoni* infection but not infected, were more likely to make errors in allotting a single yolk to a single egg case. These snails could have possessed low susceptibility (resistance) to infection by the parasite, which has been previously associated with low fecundity and abnormal egg production (Minchella & LoVerde, 1983, Cooper et al., 1994). Snails may be allocating less energy towards ensuring proper reproduction leading to the increased number of inconsistencies in egg cases (multiple yolks or no yolks). Instead, this energy could be allocated more towards immunological maintenance to limit the potential risk of infection.

In this study, we explored the consequences of *S. mansoni* infection on the reproductive success of *B. glabrata*. We not only add to the evidence for fecundity compensation, we expand the definition by showing that it can occur both by increased probability of egg laying as well as increased number of eggs produced. Despite the increase in egg production by exposed hosts, offspring of infected snails appear to do just as well as uninfected snail offspring. We additionally see evidence that resistance may have an impact on egg laying, but more work on the mechanistic underpinnings of these results is needed. Our understanding of host-parasite interactions could benefit from integrating with dynamic energy budget theory to better understand how hosts allocate energy and the optimal strategies for maximizing reproductive output while combating infection (Civitello, et al., 2018; Hall et al., 2007).

## Ethical standards

The authors assert that all procedures contributing to this work comply with the ethical standards of the relevant national and institutional guides on the care and use of laboratory animals.

## Acknowledgements

We would like to sincerely thank the members of the Minchella laboratory group, which without their assistance, completion of this study would not have been possible. B. glabrata snails provided by the NIAID Schistosomiasis Resource Center of the Biomedical Research Institute (Rockville, MD) through NIH-NIAID Contract HHSN272201700014I for distribution through BEI Resources. Annabell Davis was supported by a Cable-Silkman fellowship for undergraduates in Parasitology and the Summer Stay Scholarship provided by Purdue University.

